# Structural insights into the RNA maturation of the mitoribosome by GTPBP10

**DOI:** 10.1101/2023.11.07.565989

**Authors:** Thu Giang Nguyen, Christina Ritter, Eva Kummer

## Abstract

Mitochondria contain their own genetic information and a dedicated translation system to express it. The mitochondrial ribosome is assembled from mitochondrial-encoded RNA and nuclear-encoded ribosomal proteins. Assembly is coordinated in the mitochondrial matrix by biogenesis factors that transiently associate with the maturing particle. Here, we present a structural snapshot of a previously unknown large mitoribosomal subunit assembly intermediate containing 7 biogenesis factors including the GTPases GTPBP7 and GTPBP10. Our structure illustrates how GTPBP10 aids the folding of the ribosomal RNA during the biogenesis process, how this process is related to bacterial ribosome biogenesis, and why mitochondria require 2 biogenesis factors in contrast to only 1 in bacteria.

## Introduction

Mitochondria are eukaryotic organelles with a central role in cell metabolism, regulation of apoptosis, cell differentiation, and innate immunity. They contain their own genetic material that encodes for approximately a dozen essential components of the respiratory chain complexes in humans. To express their genome, mitochondria maintain a distinct translation apparatus that differs from its cytosolic counterpart and the bacterial ancestor. Especially, the architecture and composition of the mitochondrial ribosome (mitoribosome) has changed representing an unusually protein-rich ribosome with reduced RNA elements in humans.^1,2^ The human mitochondrial ribosome is assembled in the mitochondrial matrix from ribosomal RNA (rRNA) encoded on the mitochondrial DNA and ribosomal proteins of nuclear origin. After their syntheses in the cytosol, the mitochondrial ribosomal proteins are imported into the organelle and associate with the rRNA in a coordinated and hierarchical fashion.^3,4^

The assembly of the mitoribosomal small and large subunits (mtLSU and mtSSU, respectively) is assisted by dedicated biogenesis factors that temporarily bind to the maturing particles to establish the timely and coordinated incorporation of ribosomal proteins as well as the folding and modification of rRNA. Biogenesis factors are both, conserved or mitochondria-specific and include methyltransferases and pseudouridine synthases, helicases, GTPases, and other proteins.^3-5^ Defects in ribosome biogenesis reduce mitochondrial protein synthesis and cause severe human diseases including encephalomyopathy, optic neuropathy, and spastic paraplegia.^6^

Recently, several single particle cryo-electron microscopy (cryo-EM) structures of biogenesis intermediates of the human mtLSU have provided a detailed view of the later stages of the assembly process, where the outer shell of the particle has already formed but the subunit interface is still immature. These structure have been vital to understand the order, in which the single components are assembled into the final functional mitoribosome, as well as the positioning, function, and interplay of maturation factors in this process.^7-12^ A remarkable feature of the assembly of the mtLSU is that the catalytic site of the ribosome - the peptidyl transferase center (PTC) - folds last to ensure that only properly assembled particles enter the translation process.^7^ This quality control checkpoint is highly conserved and a common feature in the biogenesis of the ribosome in all translation systems.^3^ Despite the wealth of mtLSU maturation snapshots, a key intermediate has so far escaped structural elucidation. Here, we present the structure of this missing late-stage maturation intermediate of the human mtLSU. Our complex contains the NSUN4-MTERF4 dimer, the MALSU1-L0R8F8-mtACP module, and the GTPases GTPBP7 and GTPBP10 at the immature, mitoribosomal large subunit interface. While the NSUN4-MTERF4 dimer and the MALSU1-L0R8F8-mtACP module have been visualized in many intermediates,^8-12^ the complex of GTP binding protein 7 (GTPBP7) and 10 (GTPBP10) is shown here for the first time.

GTPBP10 (also termed OBGH2) is one of two mitochondrial homologs of the bacterial GTPase ObgE with GTPBP5 (also termed OBGH1 or MTG2) being the other one. Biochemical and cell biological work, has previously identified GTPBP5 and GTPBP10 as GTPases that bind to late mtLSU maturation intermediates.^13-16^ Loss of GTPBP10 as well as GTPBP5 leads to reduced levels of 16S rRNA, a reduction in mitochondrial protein synthesis and consequently cell growth. Both, GTPBP5 and GTPBP10 can complement for loss of ObgE in E. coli,^17^ but have nonoverlapping, essential functions in human mitoribosome biogenesis.^13-16^ Intriguingly, GTPBP10 and GTPBP5 have largely similar predicted folds and differ mostly in their Obg domains where GTPBP10 contains truncations in loop1 and loop3. How these structural differences in GTPBP10 and GTPBP5 convey distinct molecular functions during the mitoribosome assembly process remains unknown. GTPBP10 has been assigned to a low-resolution EM map previously.^10^ However, in our near-atomic EM reconstruction, we discover it in a different conformation on the mtLSU and in contact with the GTPase GTPBP7. Our data permit to rationalize its role in active site maturation via H89 positioning. Our structure also shows why the mitoribosomal maturation process requires two distinct homologs of the highly conserved bacterial maturation factor ObgE.

## Results

### Isolation of assembly intermediates of the human mitoribosomal large subunit

We purified mitoribosomes in the presence of the nonhydrolyzable nucleotide analog GMPPNP from actively growing HEK293-6E cell. We rationalized that mitochondrial biogenesis and mitoribosomal activity will be high under these conditions. The mitoribosomal pool was analyzed by single particle cryoEM. We computationally isolated the mitochondrial ribosome in complex with translation elongation factor mtEFG1, the initiating 55S mitoribosome containing mtIF2, a contamination of 80S cytoplasmic ribosomal particles with elongation factor eEF2 bound, as well as 3 distinct mtLSU maturation intermediates (Fig. 1A). The mtLSU maturation intermediates contained either the biogenesis factors NSUN4-MTERF4, and the MALSU1-L0R8F8-mtACP module in the presence of GTPBP5 (intermediate 3), NSUN4-MTERF4 and the MALSU1-L0R8F8-mtACP module in the presence of GTPBP7 (intermediate 2), or NSUN4-MTERF4 and the MALSU1-L0R8F8-mtACP module in the presence of GTPBP7 and GTPBP10 (intermediate 1) (Fig. 1A). As the high-resolution structures of most of the complexes have been described elsewhere,^8,9,12^ we decided to focus our attention on the so far unresolved maturation intermediate containing the GTPases GTPBP7 and GTPBP10.

**Fig 1.**
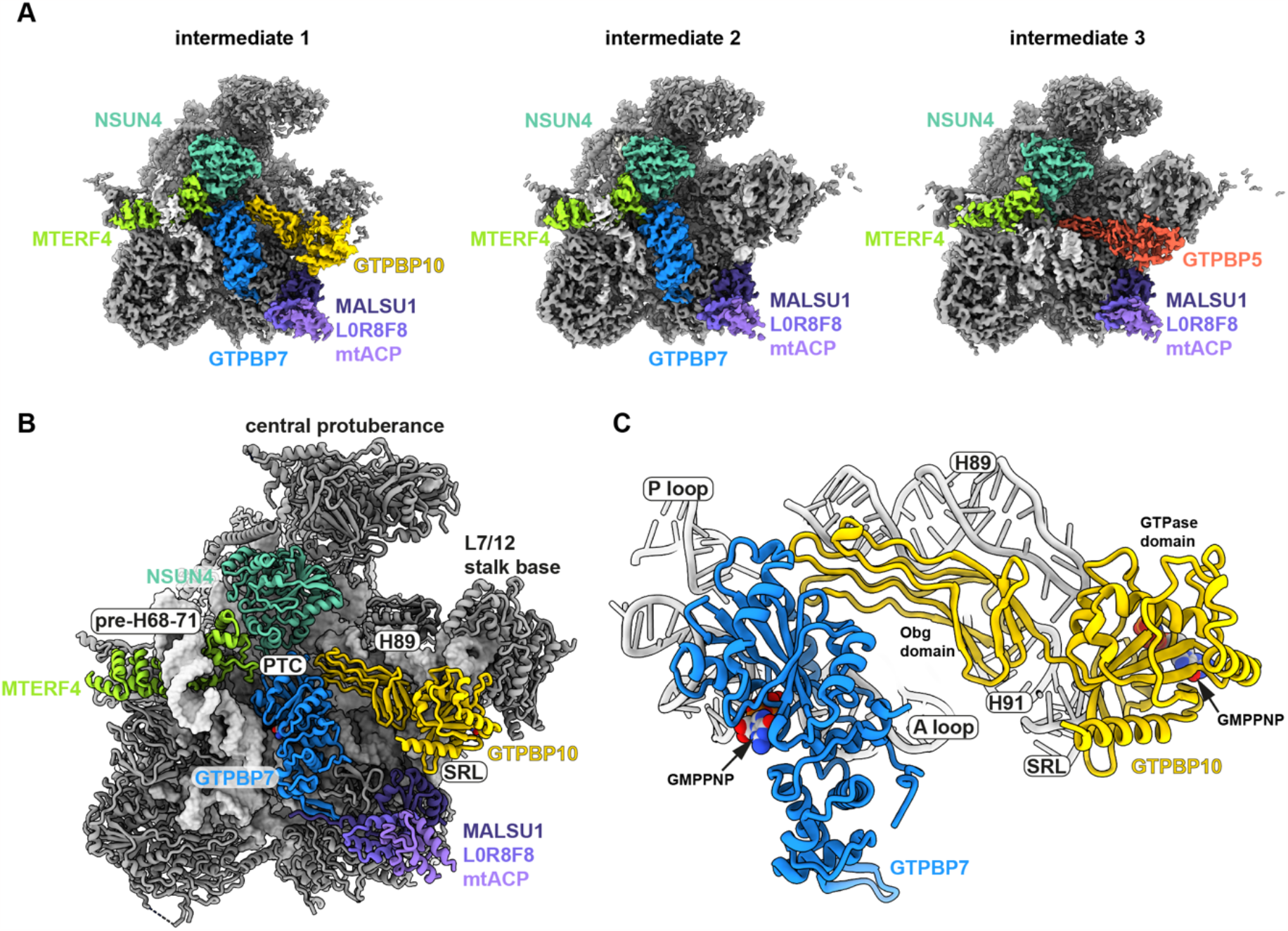
Overview of mtLSU biogenesis intermediates. **A)** Cryo-EM densities of the three isolated mtLSU maturation intermediates with the associated biogenesis factors coloured. The colours of the protein name and respective densities are matched. **B)** Overview of the structural model of intermediate 1 containing the GTPases GTPBP10 (yellow), GTPBP7 (blue), the module of MALSU1, L0R8F8, mtACP (shades of violet), and the dimer of MTERF4 (yellowgreen) and NSUN4 (cyan). The ribosomal RNA is shown as surface representation whereas all proteins are shown as cartoons. Prominent ribosomal elements are labelled including the sarcin-ricin loop (SRL), the peptidyl-transferase center (PTC), and ribosomal rRNA helix H89 (H89). **C)** GTPases GTPBP10 and GTPBP7 are shown in isolation with important ribosomal RNA elements highlighted Both GTPases contain the non-hydrolysable nucleotide analogue Guanosine-5’-[( β,γ )-imido]triphosphate (GMPPNP) in their active sites.

### A network of maturation factors at the immature large ribosomal subunit interface

The catalytic core of the mitoribosome is mostly composed of RNA elements and folds last in the maturation process.^18^ Key RNA elements encompass the PTC with A, P, and PTC loops, and the GTPase activating center (GAC) that consists of RNA and protein elements around the L7/12 stalk base including the sarcin ricin loop (SRL), and protein uL11m. While the GAC serves the binding and activation of translational GTPases, the PTC is the site of peptide bond formation where A and P loops facilitate binding of the acceptor ends of A and P site tRNAs, respectively.^19-22^

In our structure, we find the immature PTC surrounded by a heterodimer of the methyltransferase NSUN4 and the RNA-binding protein MTERF4, as well as GTPases GTPBP7 and GTPBP10 (Fig. 1B). In addition, a heterotrimer of biogenesis and anti-association factors MALSU1, L0R8F8, and mitochondrial acyl carrier protein (mtACP) interacts with the SRL, uL14m and bL19m as seen in earlier mtLSU maturation intermediates and a ribosome rescue complex.^7-12,23^ Similar to previous observations, the MTERF4-NSUN4 dimer contains its cofactor S-adenosyl-methionine (SAM) although in a position incompatible with methylation of 16S rRNA. The heterodimer binds the double stranded rRNA elements of the pre-H68-71 region as evidenced by continuous EM density that we and others can trace from H67.^8,11^ Both, NSUN4 – MTERF4 and MALSU1 – L0R8F8 – mtACP bind early in the mtLSU maturation process and persist over multiple maturation steps.

In contrast, GTPases engage with the maturing ribosome mostly at very specific stages of the maturation process. GTPBP10 is one of the two mitochondrial homologs of bacterial ObgE. It consists of an N-terminal Obg domain and a C-terminal GTPase domain (Fig. 1C). In our structure, the protein is bound in a crevice reaching from the GAC to the immature, catalytic PTC (Fig. 1B). The GTPase domain is located between the L7/12 stalk base and the SRL in a catalytically competent orientation and contains density for non-hydrolysable nucleotide analogue Guanosine-5’-[( β,γ )-imido]triphosphate (GMPPNP) in its active site. The Obg domain interacts with H89, which is trapped in a lifted-out conformation in comparison to the mature mtLSU (Suppl. Fig. 1A and B). GTPBP10 contacts the second GTPase GTPBP7 close to the PTC, where GTPBP7 in turn binds the heterodimer of NSUN4-MTERF4 (Fig. 1B and C). GTPBP7 is a homolog of bacterial biogenesis factor RbgA, which facilitates incorporation of bL36 in bacteria.^24^ It adopts a position on the maturing mtLSU earlier observed only in the absence of the methyltransferase MRM2.^12,24-27^ GTPBP7 has been proposed to monitor the 2’-*O*-ribose methylation status of the highly conserved nucleotides U3039 and possibly also G3040 in the A loop, which are critical for biogenesis and catalytic activity of the ribosome.^12,28^ In accordance with biochemical data, our maturation intermediate already contains all ribosomal proteins of the mtLSU except bL36m, whose binding site gets only accessible once H89 adopts a mature conformation.^13^

A low resolution cryo-EM structure has earlier identified GTPBP10 at the maturing mtLSU interface, but has placed it in a non-canonical conformation with the Obg instead of the GTPase domain contacting the SRL.^10^ Also, the rRNA helix H89 is in a distinct conformation in that reconstruction and displays major clashes with the putative GTPBP10. In our near-atomic resolution structure, we now unambiguously identify GTPBP10 that is in contact with other maturation factors and engages in extensive interactions with H89. It moreover associates with the catalytic center of its GTPase domain to the SRL indicating our conformation displays a catalytically competent state (Fig. 1B and C, Suppl. Fig. 2A and B). It could be possible that the earlier report shows a pre-accommodation conformation of GTPBP10 whereas our structure displays it in the final accommodated, catalytically active position.

### The role of GTPBP10 in H89 maturation

Recently, a cryo-EM structure of the homologous GTPase ObgE has been solved on the native pre-50S LSU in *E. coli*.^29^ Structure and biochemical evidence provide mechanistic insight how ObgE aids the organization of the LSU catalytic center during bacterial ribosome biogenesis. The structure highlights a role of ObgE in folding of the 23S rRNA helix H89, allowing incorporation of ribosomal proteins uL16 and bL36 in a subsequent assembly step.

In human mitochondria, two homologues of the essential ObgE exist, namely GTPBP5 (OBGH1) and GTPBP10 (OBGH2). Structural insights into GTPBP5 bound to an mtLSU intermediate indicate that H89 has already adopted an almost fully mature conformation (Suppl. Fig. 1C). In contrast, our structure shows that the GTPBP10-bound state is characterized by a large displacement of H89, as has been observed in the presence of ObgE in bacteria, indicating that GTPBP10 acts prior to GTPBP5 during mitochondrial ribosome biogenesis. While H89 is almost completely folded and most of its base pairing has been established in the GTPBP10 maturation intermediate, the H89 base is entirely disordered to allow the RNA to adopt this lifted conformation (Suppl. Fig. 1B). Moreover, the tip of H89 is stretched by clamping between the L7/12 stalk base and the Obg domain of GTPBP10 (Fig. 1B). GTPBP10 fixes H89 in this conformation mostly via electrostatic interactions of the Obg domain with the sugar-phosphate backbone of H89 (Fig. 2A). Moreover, an alpha-helical element (R52-R66), that is lacking in GTPBP5, clamps underneath the tip of rRNA helix H89 (Fig. 2B). In addition, the GTPBP10 GTPase domain is situated on the SRL and its Obg domain establishes extensive contacts to rRNA helix H91 and GTPBP7 (Fig. 1C and 2B). These interactions position GTPBP10 on the maturing particle allowing it to act as a ‘door stopper’ for H89.

**Fig 2.**
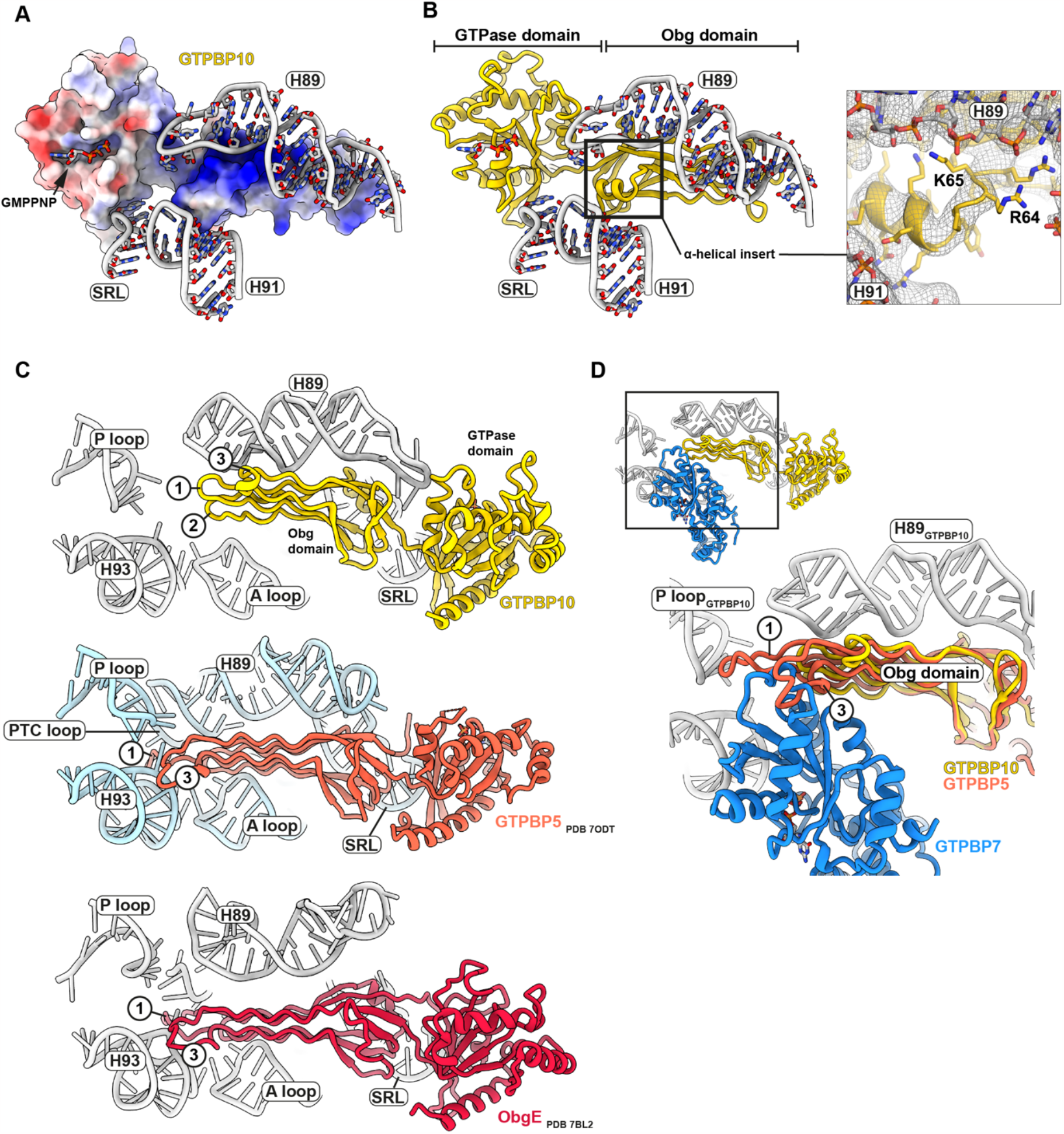
Comparison of mitochondrial GTPBP10, GTPBP5, and bacterial ObgE. **A)** Surface representation of GTPBP10 colored according to coulombic electrostatic potential using default settings (blue = positive (10), red = negative (-10)) in ChimeraX. Ribosomal RNA helices H89, H91 and the sarcin-ricin loop (SRL) are shown for reference. **B)** GTPBP10 structural model shown in the same orientation as in A). Organization and positioning of GTPase and Obg domains are highlighted and the non-hydrolysable nucleotide analog GMPPNP is shown in its binding site. Important surrounding ribosomal RNA elements are shown. The GTPBP10-specific alpha helical element that aids H89 positioning is boxed and shown in more detail on the right. The corresponding EM density has been included as mesh, gaussian-filtered at 1σ. **C)** A comparison of the mitochondrial ObgE homologs GTPBP10 and GTPBP5 with its bacterial counterpart is shown. For comparison the ribosomal RNA of the large subunit from intermediate 1, PDB 7ODT (GTPBP5), and PDB 7BL2 (ObgE) has been superimposed and important, conserved ribosomal elements are highlighted in the images. It is apparent that the proteins diverge in the length of their Obg domain loops 1 and 3, which have been labelled with numbers in the images. **D)** Superposition of the Obg domain of GTPBP5 (PDB 7ODT) onto GTPBP10 shows that binding of GTPBP5 is incompatible with the location of GTPBP7 in intermediate 1.

Overall, GTPBP10 interaction with GTPBP7, H91 and the H89 RNA backbone make it sterically impossible for H89 to slip into its final position. GTPBP10 release is necessary to resolve this steric block and allow accommodation of H89 in its binding pocket on the mtLSU. In our structure, the L7/12 stalk base is shifted outwards. H89 accommodation will allow relocation of the stalk base and uL11m. uL11m will subsequently collide with a GTPBP10-specific insertion in its GTPase domain (residues 240-247) indicating that H89 accommodation is necessarily connected to release of GTPBP10 from the GAC (Fig. 3D). The dislocation of the stalk base leaves the binding site for bL36 inaccessible in intermediate 1, in accordance with biochemical data.^13^ Although we can identify density for uL16, it is still rather loosely attached to the immature H89. Overall, our data confirm that H89 folding and accommodation via GTPBP10 are necessary for sequential incorporation of uL16 and later also bL36 into the mtLSU.

**Fig 3.**
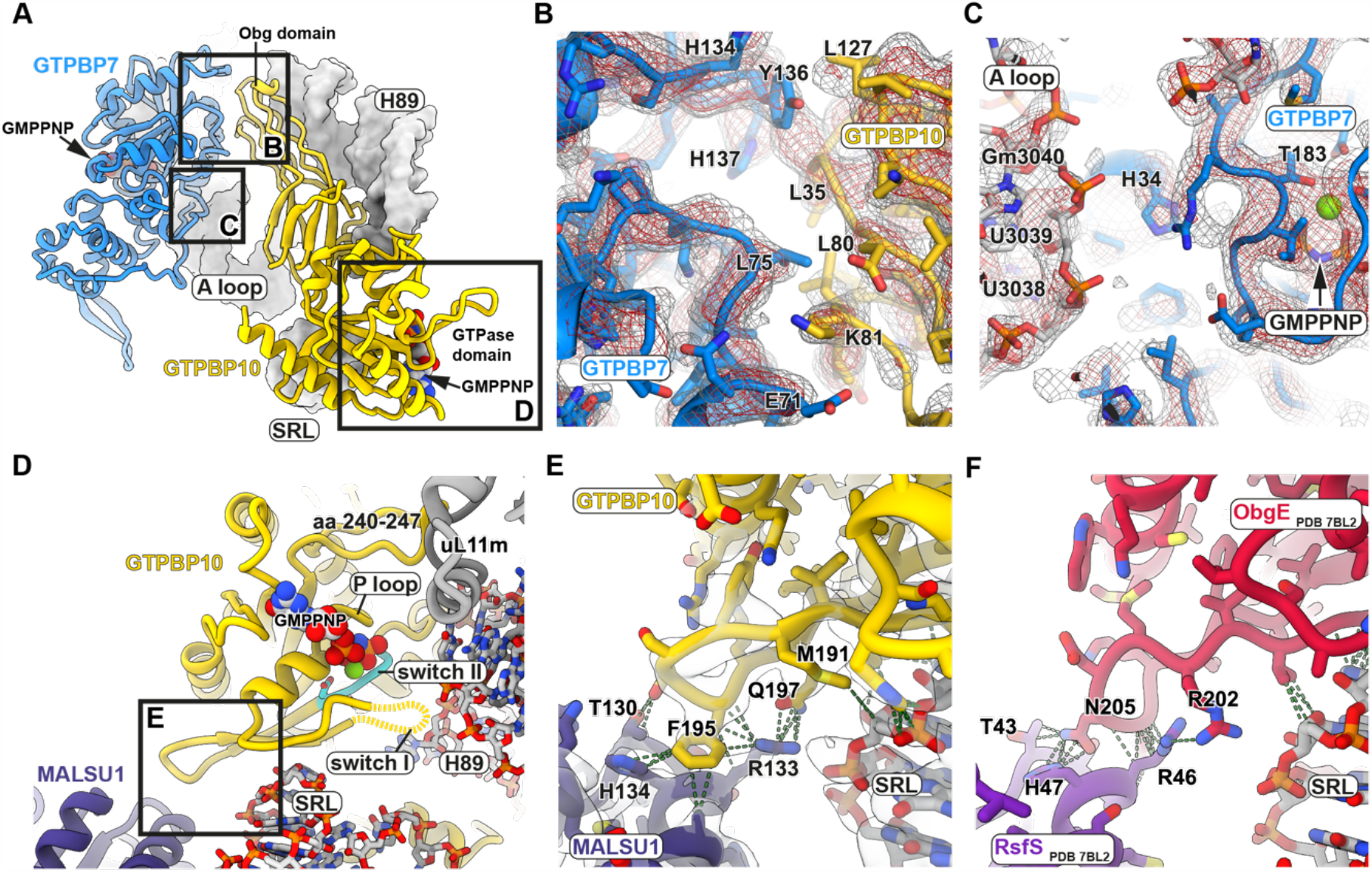
Interactions of GTPBP10 and GTPBP7 with the ribosomal RNA and other maturation factors. **A)** Overview of GTPBP7 and GTPBP10 in the context of nearby mitoribosomal RNA elements. Regions highlighted in panle B, C, and D are boxed. **B)** The mostly hydrophobic interaction interface of GTPBP10 and GTPBP7 is shown. The corresponding density is displayed as mesh at two different thresholds (red = 3.5σ, grey = 2.1σ). **C)** The location of histidine 34 (H34) of GTPBP7 in relation to the ribosomal A loop is depicted. EM density is shown at 2 different thresholds as in B). **D)** Overview of the location of the GTPase domain and the bound nucleotide in the context of ribosomal RNA helix H89, the ribosomal stalk base containing uL11m, the sarcin-ricin loop (SRL), and biogenesis factor MALSU1. Switch II containing the catalytic walker B motif (DxxG) is coloured in cyan with residue D202 highlighted. The region detailed in E is boxed. **E and F)** Contacts between GTPBP10 and MALSU1 as well as the bacterial ObgE with RsfS (PDB 7BL2) are represented as dotted lines. The EM density for intermediate 1 is displayed as semi-transparent surface. The loop likely aids to anchor the GTPBP10 GTPase domain via MALSU1 on SRL as it was one of the best-resolved regions within the GTPase domain.

### Comparison of GTPBP10 to mitochondrial GTPBP5 and bacterial ObgE

GTPBP5 and GTPBP10 have nonoverlapping, essential functions in human mitoribosome biogenesis.^13-16^ Both proteins have largely similar folds but display differences in their Obg domains (Fig. 2C). Our structure now shows that GTPBP10 contains truncations especially in loop 1 and loop 3, and an alpha-helical insertion between beta strands 2 and 3 of the Obg domain. These structural differences explain the distinct roles of both GTPases as the extended loops of GTPBP5 are sterically incompatible with GTPBP7 interaction (Fig. 2D). In contrast, the shortened loops as well as the alpha-helical element allow GTPBP10 to intimately nestle onto GTPBP7 and to interact extensively with H89 to trap it in a lifted conformation (Fig. 2B and D). The extensive interaction of the Obg domain with the RNA backbone is also reflected in a higher local density of positively charged amino acid side chains in GTPBP10 in comparison to GTPBP5 (Suppl. Fig. 3A and B). However, the extended loops of GTPBP5 allow it to reach the unstructured base of H89 to trigger its folding (Fig. 2C and D, Suppl. Fig. 3D and E). Curiously, bacterial ObgE is a chimera of both mitochondrial versions, as it contains elongated loops 1 and 3 as well as the alpha-helical element in its Obg domain explaining why it can complete both, accommodation and folding of H89 (Fig. 2C, Suppl. Fig. 3C). Overall, our data rationalize how sequential action of GTPBP10 and GTPBP5 replaces the function of bacterial ObgE in mitochondrial ribosome biogenesis.

### Interactions of GTPBP7 and GTPBP10 with rRNA elements, ribosomal proteins and each other

Our data indicate that GTPBP10 action requires the simultaneous presence of GTPBP7 on the ribosome. While the tip of H89 is stabilized via interaction of GTPBP10 with the SRL and H91, the Obg domain stacks onto GTPBP7. The interfaces of both proteins show high shape complementarity and establish the interaction mostly via hydrophobic contacts (Fig. 3A and B). In addition, GTPBP7 makes extensive contact to ribosomal RNA including the A loop, H93, and H71. GTPBP7 was proposed to monitor methylation status of U3039, G3040, and G2815 in A and P loop, respectively.^12^ In our reconstruction, GTPBP7 is too far from the P loop for a direct interaction but touches the backbone of the A loop whose terminal tip is disordered (nucleotides 3041-3043) (Fig. 3C). Our density does however not provide an indication for stacking of U3039 with His34 of GTPBP7 as proposed earlier.^12^ The 2’*O*-methyl group of the ribose of U3039 may instead directly interact with amino acid stretch Pro32/Gly33/His34 of GTPBP7 in our maturation intermediate although the resolution of our EM map does not permit to unambiguously identify atomic details of this putative interaction. His34 instead faces towards the gamma-phosphate of the bound nucleotide. We also find GTPBP7 to wrap around ribosomal protein uL14m and to bind with a beta-hairpin (residues 280-295) in a cleft between ribosomal protein uL14m and bL19m (Suppl. Fig. 3F). The GTPBP7 beta hairpin is additionally stabilized by the C-terminus of MALSU1 reaching over (Suppl. Fig. 3F). The beta hairpin has earlier been shown to serve as an anchor point around which GTPBP7 swings from a conformation contacting the ribosomal immature PTC towards pre_h68-71 upon binding of the methyltransferase MRM2 to the A loop.^11,30^

Besides its interaction with GTPBP7, GTPBP10 also contacts the biogenesis factor MALSU1 in intermediate 1. MALSU1 is a homolog of bacterial RsfS, which was proposed to cooperate with bacterial ObgE during H89 maturation.^29^ Previous studies on GTPBP5 rationalized that, distinct to the bacterial system, it acts in concert with the N-terminal tail of biogenesis factors NSUN4 instead of MALSU1.^9^ In contrast, our structural data now indicate that MALSU1 likely plays a role for GTPBP10 action as it engages in similar interactions with the GTPase as the bacterial RsfS (Fig. 3D-F) and we do not find the N-terminal tail of NSUN4 to be close to GTPBP10. The interaction with MALSU1 could aid to position the nearby catalytic walker B motif in switch II of the GTPBP10 GTPase domain on the SRL (Fig. 3D).

Our near-atomic reconstruction of GTPBP10 together with GTPBP7 now allows to rationalize previously described mutations reported to ablate GTPBP10 function in mitoribosome biogenesis.^13^ We find the mutations to potentially either disturb the fold of the Obg domain and to hamper interaction with GTPBP7 (G82E), to influence binding of the nucleotide to the GTPase domains (S325P), or to influence the interaction of GTPBP10 with 16S rRNA (deletion of R64 and K65) (Fig. 2B and Suppl. Fig. 4).^13^

## Discussion

We isolated mitoribosomal complexes from mitochondria of actively growing HEK293-6E cells in the presence of the non-hydrolysable nucleotide analogue GMPPNP. Among the isolated complexes, we discovered a previously unresolved, native biogenesis intermediate of the human mitoribosomal large subunit. Our intermediate contains 7 biogenesis factors including the heterodimer of NSUN4 and MTERF4, the MALSU1 – L0R8F8 – mtACP complex, and the two GTPases GTPBP7 and GTPBP10.

GTPBP10 is next to GTPBP5 one of two mitochondrial homologues of the bacterial GTPase ObgE. In bacteria, ObgE coordinates the folding of rRNA helix H89 and the incorporation of ribosomal proteins uL16 and bL36 to form a fully functional catalytic center.^29^ Mitochondrial ribosome biogenesis requires two distinct homologs of the bacterial GTPase to generate functional ribosomes. Together with previously published structures of the methyltransferase MRM2 and GTPBP5, our data substantiate that the two mitochondrial homologues exert complementary but distinct functions in H89 folding. The structural data also allow us to derive a more complete picture of the order of events in mtLSU biogenesis (Fig. 4).^8,9,11^ Assembly of the PTC requires coordinated action of methyltransferases and GTPases to modify and fold the ribosomal RNA. In an earlier step, the methyltransferase MRM3 catalyzes the 2’-O-methylation of G3040 in the ribosomal A loop. After MRM3 is released, the NSUN4-MTERF4 dimer associates and leads to a rearrangement of the pre-H68-71 rRNA stretch. Then, GTPBP10 catalyzes in the presence of GTPBP7 the deposition of the already largely folded H89 into its ribosomal cavity while the base of the rRNA helix remains partially unfolded. Upon H89 deposition, rearrangement of the L7/12 stalk base may lead to GTP hydrolysis in GTPBP10 causing its dissociation from the ribosomal particle. In addition, H89 motion allows uL16m and bL36m to get firmly incorporated, which may trigger GTPBP7 GTPase activity in analogy to the bacterial system, where incorporation of bL16 was shown to stimulate GTPase activity of the bacterial GTPBP7 homolog RbgA.^31^ This likely leads to a reorganization of GTPBP7 on the subunit interface around its hinge point associated to uL14m and bL19m. Now, the methyltransferase MRM2 can bind close to the PTC to catalyze 2’-O-methylation of U3039 in the A loop.^11^ Finally, GTPBP5 binds to the maturation intermediate and triggers release of the A loop from the binding pocket in MRM2 and folding of the unstructured H89 base into its final conformation in concert with the N-terminal tail of NSUN4.^8,9,11^

**Fig 4.**
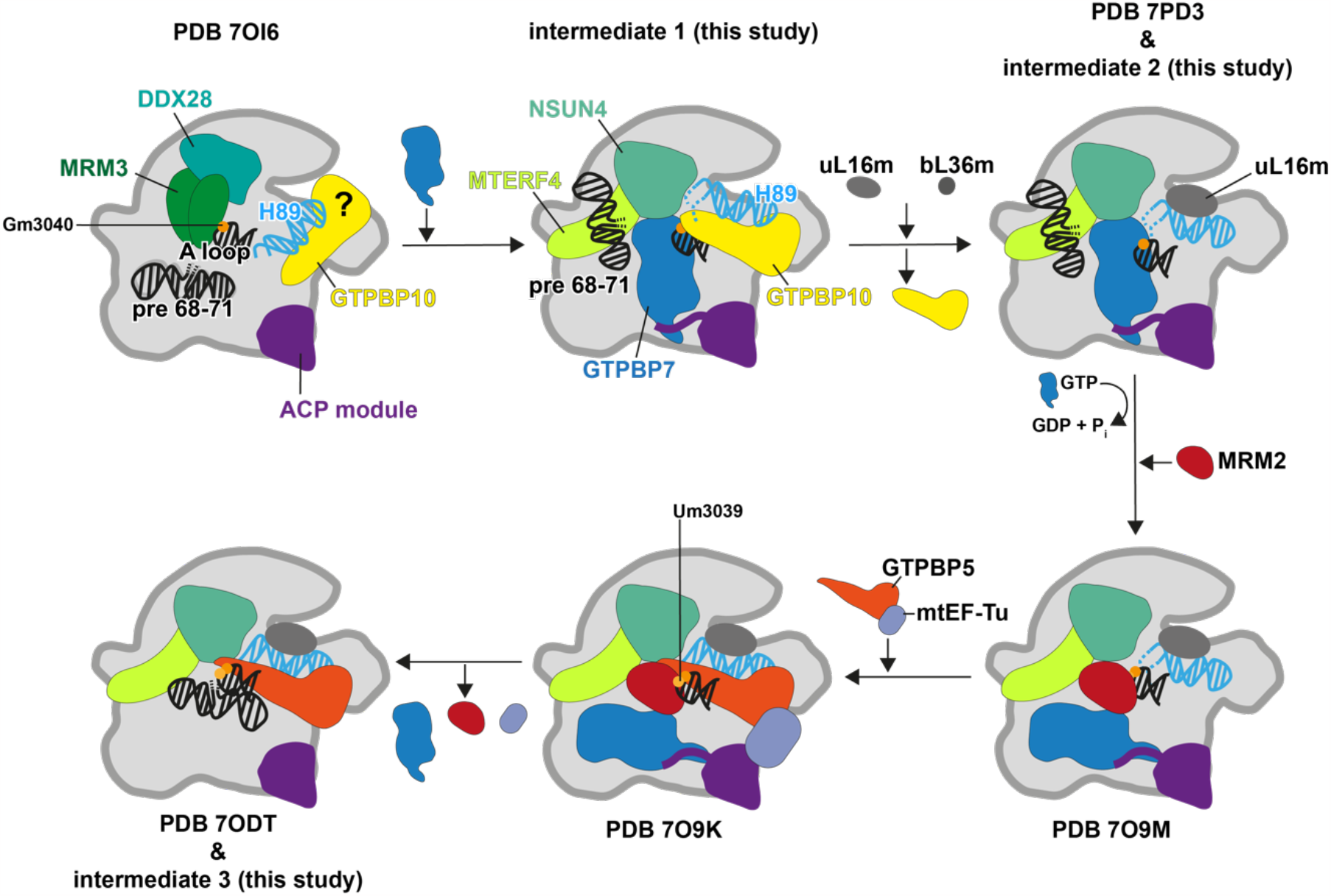
Model for H89 maturation. H89 maturation requires the concerted action of GTPBP10 and GTPBP5. GTPBP10 has earlier been proposed to be part of a maturation intermediate in conjunction with the methyltransferase MRM3 and helicase DDX28. However, the low resolution of the reconstruction renders this interpretation ambiguous. We now find GTPBP10 in concert with GTPBP7 bound to the maturing ribosome when the pre-H68-71 rRNA stretch has already been deposited on the NSUN4-MTERF4 dimer. Deposition of H89 into its ribosomal cavity enables proper incorporation of uL16m and bL36m, which in turn may trigger reorganization of GTPBP7 on the RNA surface. This will vacate the necessary space for binding of the methyltransferase MRM2, which installs the methylation on U3039 in the A loop. As shown earlier, binding of GTPBP5 release the A loop from MRM2 and may trigger departure of MRM2 and GTPBP7 from the complex.

This division of tasks between GTPBP10 and GTPBP5 can be explained by structural features.

GTPBP10 and GTPBP5 show largely similar folds but differ in their loop1 and 3 regions as well as by peptide insertions in their Obg and GTPase domains. GTPBP5 contains longer loops 1 and 3, which contact immature rRNA elements of the catalytic center. Here, they coordinate the repositioning of the P loop and the folding of the base of H89 in conjunction with the N-terminal tail of NSUN4. Moreover, GTPBP5 was found to interact with the translation elongation factor mtEF-Tu, where mtEF-TU was proposed to aid accommodation of GTPBP5 onto the mtLSU.^11^ In contrast, GTPBP10 contains shorter loops 1 and 3 of the Obg domain and an alpha helical insertion that permit tight interaction of GTPBP10 with GTPBP7 and H89, respectively. The GTPBP10 loops do not primarily engage in protein-nucleic acid interaction at the PTC and the N-terminal tail of NSUN4 is disordered in our intermediate 1. Instead, the loops have rather been repurposed to form an interaction interface with GTPBP7 to stabilize GTPBP10 binding on the rRNA. One side of the Obg domain is enriched in positively charged amino acids enabling the tight interaction with the RNA backbone of H89 and its positioning on top of its binding cavity in the mtLSU. We do not find mtEF-Tu in our intermediate 1 assembly suggesting that GTPBP10 may be able to bind to the maturing ribosomal subunit by itself, possibly due to the additional contact to GTPBP7. Collectively, the structural data explain why GTPBP10 and GTPBP5 are both essential in mitoribosome biogenesis as only their concerted action completes H89 folding.

## Materials and Methods

### Cell culture

The human embryonic kidney 293EBNA1-6E cell line (HEK 293EBNA1-6E) was adapted to growth in suspension in F17 medium (FreeStyleTM F17 Expression Medium, Gibco), supplemented with 4 mM L-Glutamine (Sigma), 0.1% Pluronic F-68 (Gibco), and 1% heat-inactivated fetal bovine serum (Sigma). The cell culture was maintained 0.8-1.0 × 10^6^ cells/ml in 1L square polycarbonate storage bottles (Corning), and shaken in a humidified incubator (Infors HT) at 150 rpm and 37 °C with 5% CO_2_. Cell density and viability were determined via the trypan blue exclusion method with the CountessTM automated cell counter (Invitrogen). Polyethylenimine (PEI) was used as a transfection reagent for transient overexpression of mtRF1-AAG-3xFLAG in HEK 293EBNA1-6E cells, following a previously described method with some modifications.^33^ Cell culture with viability greater than 98% was split 3 hours before transfection in fresh complete F17 medium to 1.0 × 10^6^ cells/ml. Plasmid DNA of mtRF1-AAG-3xFLAG in a pcDNA3.1 vector backbone was ordered from Thermo Scientific. DNA and PEI at a ratio of 1:3 (w/w) were diluted in F17 medium to make up a transfection reagent volume as 1% of the culture volume, with final DNA concentration as 1 mg per liter of culture. The transfection mixture was incubated 20 min at room temperature to form polyplexes before being added to the culture. 2 days after transfection, cells were harvested and used for the next steps.

### Mitochondria isolation

1L of HEK293-EBNA1-6E cell culture at a density of 1.5 - 2.0 × 10^6^ cells/mL was harvested by centrifugation (2000 rpm, 4 °C, 15 min) using a Sorvall SLC-4000 rotor (Thermo Fisher). Cells were washed in 20 mL of chilled phosphate-buffered saline (1X PBS, pH 7.4), and centrifuged (1500 ×g, 4 °C, 15 min) using the Centrifuge 5810R (Eppendorf). The cell pellet was subsequently resuspended in 30 mL of ice-cold RSB hypo buffer (10 mM NaCl, 1.5 mM MgCl2, 10 mM Tris-HCl, pH 7.5) to allow cells to swell. After a 10 min incubation, swollen cells were broken by a Dounce homogenizer 100 mL tube and a B (tight) pestle with 15 strokes. The Dounce homogenizer was chilled on ice beforehand, and filled with a volume of 2.5X MS homogenization buffer (525 mM mannitol, 175 mM sucrose, 2.5 mM EDTA, 2.5 mM DTT, 12.5 mM Tris-HCl, pH 7.5) that was calculated accordingly to obtain a final concentration of 1X MS homogenization buffer after adding the cell resuspension.

Mitochondria was isolated from the above homogenate by differential centrifugation. The supernatant containing mitoplasts was carefully collected after each round of centrifugation at 1300 ×g then 3000 ×g (4 °C, 15 min, Centrifuge 5810R). Mitochondrial crude was pelleted by centrifugation at 9500 ×g (4 °C, 15 min) using the Optima XE-90 Ultracentrifuge with a Ti-45 rotor (Beckman Coulter). The final pellet was resuspended in 2 mL of Resuspension buffer (250 mM sucrose, 1 mM EDTA, 20 mM HEPES-KOH, pH 7.6), snap-frozen in liquid nitrogen and stored at -80 °C.

### Mitoribosome isolation

1 mL of mitochondria suspension (corresponding to 0.5 L HEK293-EBNA1-6E suspension cell culture at 1.5 - 2.0 × 10^6^ cells/mL) were supplemented with 100 uL of 100 mM GMPPNP (Jena Bioscience, NU-899-50) and quickly thawn in a water bath at room temperature. The mitochondrial suspension was mixed with 1.75 mL of lysis buffer (20 mM HEPES-KOH pH 7.6, 100 mM KCl, 40 mM MgCl2, 40 U/mL Ribolock RNase inhibitor (Thermo Scientific, 11581505), 1 tablet / 50 mL of Pierce Protease Inhibitors without EDTA (Thermo Scientific, 15677308), 0.8 mM spermidine pH 7.5, 1 mM DTT). 750 uL of 6x solubilisation buffer (20 mM HEPES-KOH pH 7.6, 100 mM KCl, 40 mM MgCl2, 9.6 % (v/v) Triton X-100 (VWR, M143-1L), 1 mM DTT) were added and the sample was gently mixed by inversion. The final volume was around 3.5 mL in a 15 mL Falcon tube. The suspension was placed on the rotation wheel for 20 min in the cold room for gentle agitation. Then the suspension was cleared 2 times for 10 min at 21300 rcf and 4 °C. The supernatant was transferred into a fresh 15 mL Falcon tube. 1 open-top, thinwall, ultraclear ultracentrifugation tube (Beckman Coulter, 344057) was filled with 1.3 mL of 40 % sucrose solution (20 mM HEPES-KOH pH 7.6, 100 mM KCl, 40 mM MgCl2, 40 % (w/v) sucrose, 1 mM DTT). 3.2 mL of the cleared, mitochondrial lysate was then carefully placed onto the sucrose cushion (ration of cushion:lysate was 1:2.5). The sample was spun for 4 h at 45.000 rpm at 4 °C in a SW55 rotor using a Beckman Coulter Optima XE-90 ultracentrifuge. The supernatant was taken off and the pellets was rinsed 2 times with 500 uL of 1x monosome buffer (20 mM HEPES-KOH pH 7.6, 100 mM KCl, 40 mM MgCl2, 1 mM DTT). The rinse was discarded and the pellets were submerged in 100 uL of 1x monosome buffer containing 1 mM GMPPNP. The pellets were resuspended by gentle shaking in the cold room for approx. 1 h. The remaining pellet was gently resuspended with the pipette and the ribosomal suspension was transferred into a 1.5 mL eppi. The suspension was cleared 2 times at 21300 rcf for 10 min at 4 °C. The supernatant was tested in negative stain to verify the ribosome concentration, which was estimated to be around 70 nM. The suspension was used directly for cryo-grid preparation.

### Cryo-EM sample preparation and data collection

Quantifoil R 1.2/1.3 or R 2/2 plus C2 on 300 copper mesh were glow discharged for 30 sec at 5 mA in a Leica Coater ACE 200. 5 uL of sample were applied to the grids and the sample was vitrified using a FEI Vitrobot Mark IV at 100 % humidity, blot force 0, wait time of 30 sec, and blotting times between 3 and 7 sec. 2 data sets were collected as movies at a pixel size of 1.08 Å/px, 300 kV and 40 e/Å2 with 33 or 42 fractions, respectively, on a FEI Titan Krios G2 equipped with Falcon 3 DED using EPU–Automated Acquisition Software (v2.14.0).

### Cryo-EM data analysis

Movies were aligned, dose weighted, and summed into micrographs using patch motion correction and patch CTF in CryoSPARC v.4.2.0.^34^ Particles were identified using the blob picker tool with a minimum and maximum particle diameter of 150 Å and 350 Å, respectively. Afterwards picks were further filtered adjusting NCC score, as well as lower and upper local power thresholds to minimize false positive picks.

For data set 1, 8’901’726 particles were extracted from 35’399 micrographs with a box size of 480 px and Fourier cropped to 120 px. Particle images underwent 2D classification in batches for at least 60 online EM iterations. Well-resolved 2D classes were 2D classified one more time to remove remaining poor particle images from the image pool. The poor particle classes from the first 2D classification also underwent one more round of 2D classification to retrieve any good remaining particle images. A total of 2’764’332 particle images were finally selected for further 3D hetero refinement in 3 batches using reference volumes including 39S mtLSU volumes, 55S mitoribosome volumes, 28S mtSSU volumes, and 80S cytosolic ribosome volumes. Reference volumes were obtained via subset selection of 2D classes resembling the respective particle type and ab initio reconstruction with 2 classes. The ab initio class that was better resolved was then used as the reference volume.

At this point, it became obvious that 39S volumes showed partially inhomogeneous subunit interfaces hinting at the presence of 39S maturation intermediates in our ribosome preparation. To obtain these putative 39S maturation intermediates, the particle image subsets belonging to well-defined 39S 3D volumes were pooled, subjected to 3D homogeneous refinement, and further classified via local 3D classification in 10 classes without resolution restriction and without image alignment using a mask covering the 39S subunit interface. Classes that contained clear densities for maturation factors were pooled yielding 125’003 particle images. Particle images were re-extracted at full pixel size and a box size of 480 px. They were local 3D classified into 4 classes with a mask surrounding the GTPase binding site between GAC and PTC. 1 class represented the mtLSU containing GTPBP7 and GTPB10, one class contained only GTPBP5, and two classes contained only GTPBP7. The class containing BP7 and 10 with 29510 particle images was homogeneously refined and further local 3D classified using the same mask as in the step before to clean the particle population from false positive images that did not contain GTPBP10 yielding 17’274 good particle images, which were joined with the final GTPBP10 particle subset of 14’382 particle images from dataset 2 for homogenous refinement, local CTF estimation, and a final round of homogenous refinement resulting in a 3D reconstruction of 3.15 Å from 31’656 particle images.

For data set 2, 8’005’604 particle images were extracted from 30’287 micrographs with a box sixe of 480 px and Fourier cropped to 120 px. Particle images underwent a similar classification procedure as for data set 1 with minor modifications (see classification scheme in Suppl. Fig. 5 for intermediate 1).

In addition to intermediate 1, we also deposited the EM maps for intermediate 2 and 3 but did not perform model building on them as the structures have been published earlier. Classification schemes for intermediate 2 and 3 are provided in Suppl. Fig. 6 and 7, respectively.

### Model building

We used the published LSU maturation intermediate containing GTPBP5 (PDB: 7ODT)^8^ and an AlphaFold model of GTPBP10 (referring to UniProt ID A4D1E9) as starting point for model building. RNA elements that were not visible in our EM map were removed in Coot.^35,36^ The Obg and GTPase domains of GTPBP10 were fitted separately as rigid bodies in UCSF Chimera.^37^ Afterwards, we revised the model manually in Coot. The RNA double helix of H89 was fitted as rigid body into its respective density and manually adjusted to account for changes especially in the tip and base region. All maturation factors as well as all ribosomal protein and ribosomal RNA were adjusted to account for the EM density either via manual model refinement or rigid body fitting of secondary structure elements and domains depending on the resolution of the respective region of the EM map. Non-hydrolyzable nucleotide analog GMPNP was added to GTPBP7 and GTPBP10 and coordinated with a Mg2+ ion and the respective residues in the active site of both GTPase. The final model was realspace-refined in Phenix using defaults restraints (Ramachandran, C-beta deviations, rotamer, secondary structure) and global minimization as well as B factor refinement in 5 cycles with a weight of experimental data and restraints set to 1.5.^38^ FSCs of the half sets of the experimental data for intermediates 1, 2, and 3 as well as of the full map with the structural model for intermediate 1 have been calculated in PHENIX using *phenix*.*mtriage*.^39^ Additonally, local resolution estimations have been carried out in crypSPARC v.4.2.0. All data including angular distribution of particle images are shown for intermediate 1 (Suppl. Fig. 8A), intermediate 2 (Suppl. Fig. 8B), and intermediate 3 (Suppl. Fig. 8C).

### Figure preparation

Molecular graphics were generated using the PyMOL (Schroedinger), UCSF Chimera or ChimeraX packages.^37,40^

## Data availability

Electron microscopy data have been deposited in the EMDB under accession codes EMD-XXXXX (intermediate1), EMD-XXXXX (intermediate2), and EMD-XXXXX (intermediate 3). The structural model for intermediate has been deposited in the PDB database under the accession code XXXX.

## Supporting information

SupplementaryInformation_Nguyen2023

## Acknowledgements

We would like to thank the staff at the Core Facility for Integrated Microscopy, Faculty of Health and Medical Sciences, University of Copenhagen for their support during EM data collection. Moreover, we would like to acknowledge Søren Kirk Amstrup for initial efforts in the model building process. This work has been supported by a Hallas-Møller Emerging Investigator grant from the Novo Nordisk Foundation to E. Kummer (NNF21OC0067360).

## Author contributions

EK designed the study and experiments. TGN carried out the cell culture work and isolated mitochondria. TGN and EK isolated mitoribosomes and vitrified samples. EK collected and analyzed EM data. EK, CR, and TGN build and interpreted the structural model. EK, CR, and TGN wrote the manuscript and prepared the figures.

