## SupplementaryInformation_Nguyen2023 for "Structural insights into the RNA maturation of the mitoribosome by GTPBP10"

### Supplementary Figures

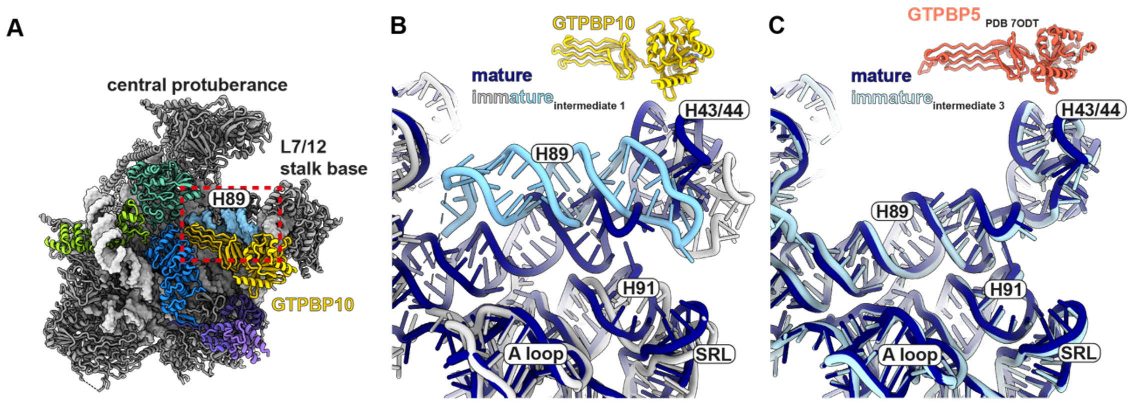


**Suppl. Fig.1 H89 displacement in maturation intermediate 1**

**A)** An overview of the structural model of intermediate 1 is shown with rRNA helix H89 highlighted in sky blue. GTPBP10 (yellow) extensively contacts the helix to stabilize it in a position on top of its crevice on the large mitoribosomal subunit. **B)** Ribosomal RNA from the immature intermediate 1 (light grey) and the mature mitoribosome (PDB 7NSH, dark blue)[32] have been superimposed to illustrate that many ribosomal RNA elements have already adopted their final location in intermediate 1 except helix H89 (sky blue). **C)** Superposition of the same ribosomal RNA elements as in A) for the maturation intermediate containing the second mitochondrial ObgE homolog GTPBP5 (PDB 7ODT, light blue). In this case, H89 is already fully accommodated.

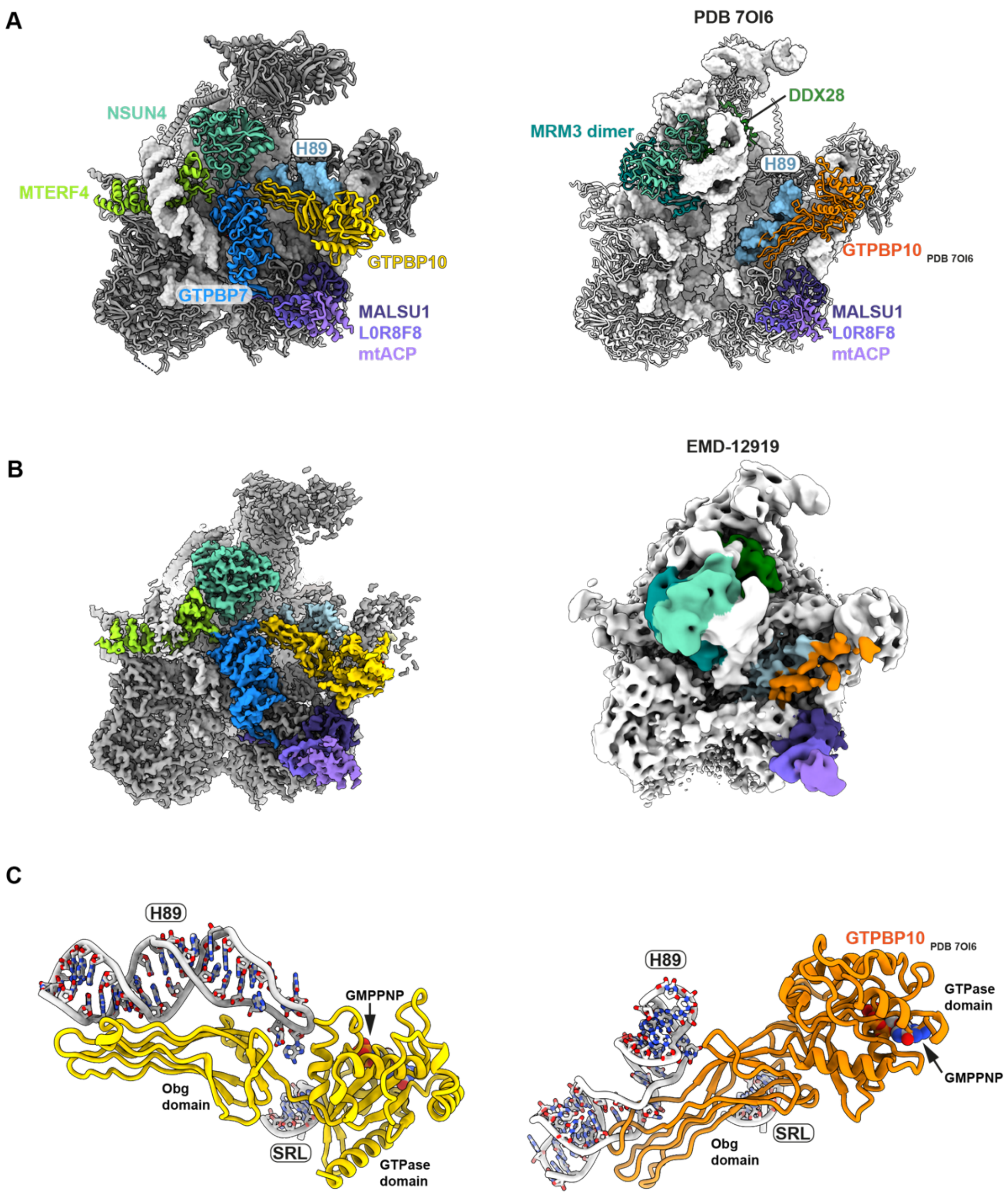


**Suppl. Fig.2 Comparison of intermediate 1 with a previously published intermediate.**
**A)** Overview of GTPBP10-containing intermediate 1 and the maturation intermediate previously published with the putative GTPBP10 (PDB 7OI6)[10]. The previous publication assigns GTPBP10 to adopt a distinct conformation on the large ribosomal subunit that is incompatible with GTP hydrolysis and occurs in the absence of maturation factor GTPBP7 at a distinct step in the maturation process. Moreover, no contact to MALSU1 and the GTPBP10 GTPase domain can be seen. **B)** Experimental EM maps for the complexes displayed in A) color-coded according to the corresponding PDBs. The experimental density for intermediate 1 of this study is shown on the left. The experimental map from PDB 7OI6[10] is shown on the right. **C)** Detailed view of A) demonstrating a different interaction of the GTPase domain with the sarcin ricin loop (SRL) and major clashes of GTPBP10 with RNA helix H89 in PDB 7OI6.

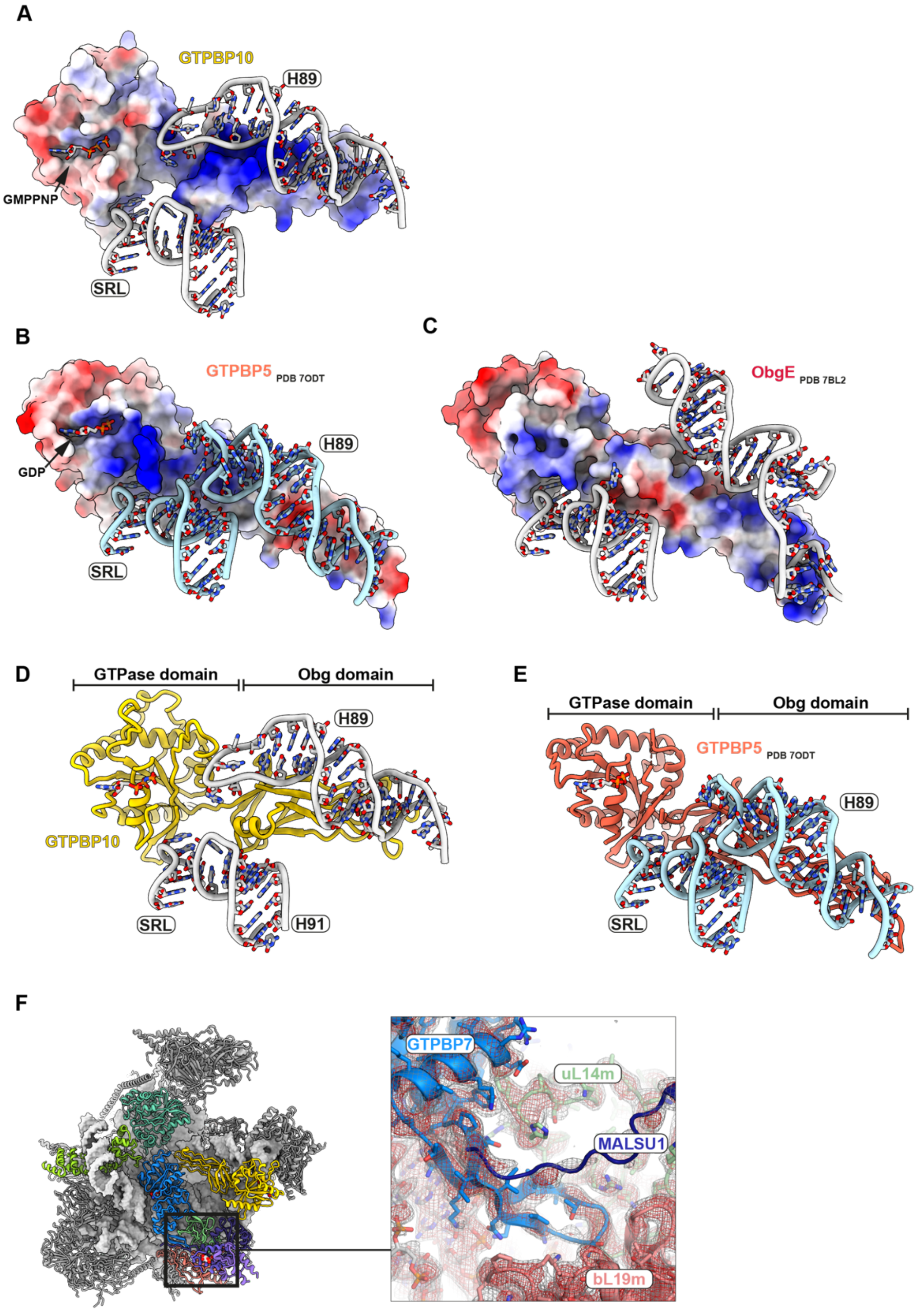


**Suppl. Fig.3 Comparison of GTPBP10, GTPBP5 and bacterial ObgE**

**A-C)** Surface representations of the Obg proteins coloured according to coulombic electrostatic potential using default settings (blue = positive (10), red = negative (-10)) in ChimeraX. Structural models for GTPBP5 and surrounding RNA elements were derived from PDB 7ODT, and for ObgE from PDB 7BL2. Surrounding RNA elements including the sarcin-ricin loop (SRL), RNA helix H89, and RNA helix H91 are shown as cartoons. **D-E)** Structural models for GTPBP10 and GTPBP5 shown in the context of important ribosomal RNA elements. The different position of H89 is apparent as well as the absence of the alpha-helical insertion in the Obg domain of GTPBP5. F) Overview of the interaction of GTPBP7 with ribosomal elements on the left. The region boxed in the overview is detailed on the right. A beta-hairpin of GTPBP7 serves to anchor the protein between ribosomal proteins uL14m and bL19m. In addition, the C-terminal tail of MALSU1 reaches over to contact GTPBP7. The EM density is shown at 2 thresholds (red = 3.5$\sigma$, grey = 2.1$\sigma$).

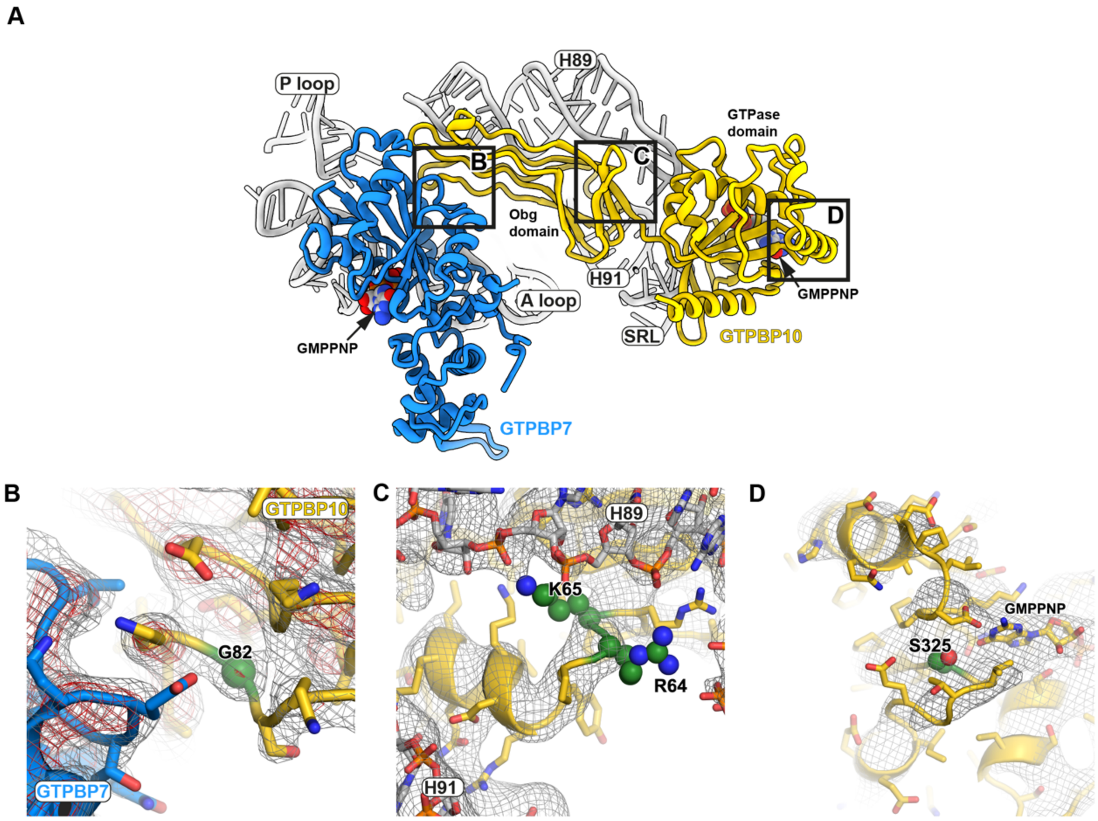


**Suppl. Fig.4 Mutations disturbing GTPBP10 function in mitoribosome biogenesis**
**A)** An overview of GTPBP10 and GTPBP7 in the context of ribosomal RNA is shown and the regions, in which the mutations are located are highlighted with boxes corresponding to the panels shown below. **B-D)** Mutations are highlighted in green and as spheres in the structural model of GTPBP10 with the EM density included. The density is shown for the sharpened map in B) at 2 thresholds (red = 3.5$\sigma$, grey = 2.1$\sigma$), and for the gaussian-filtered (1$\sigma$) map at 1 threshold (2.8$\sigma$) in C) and D).

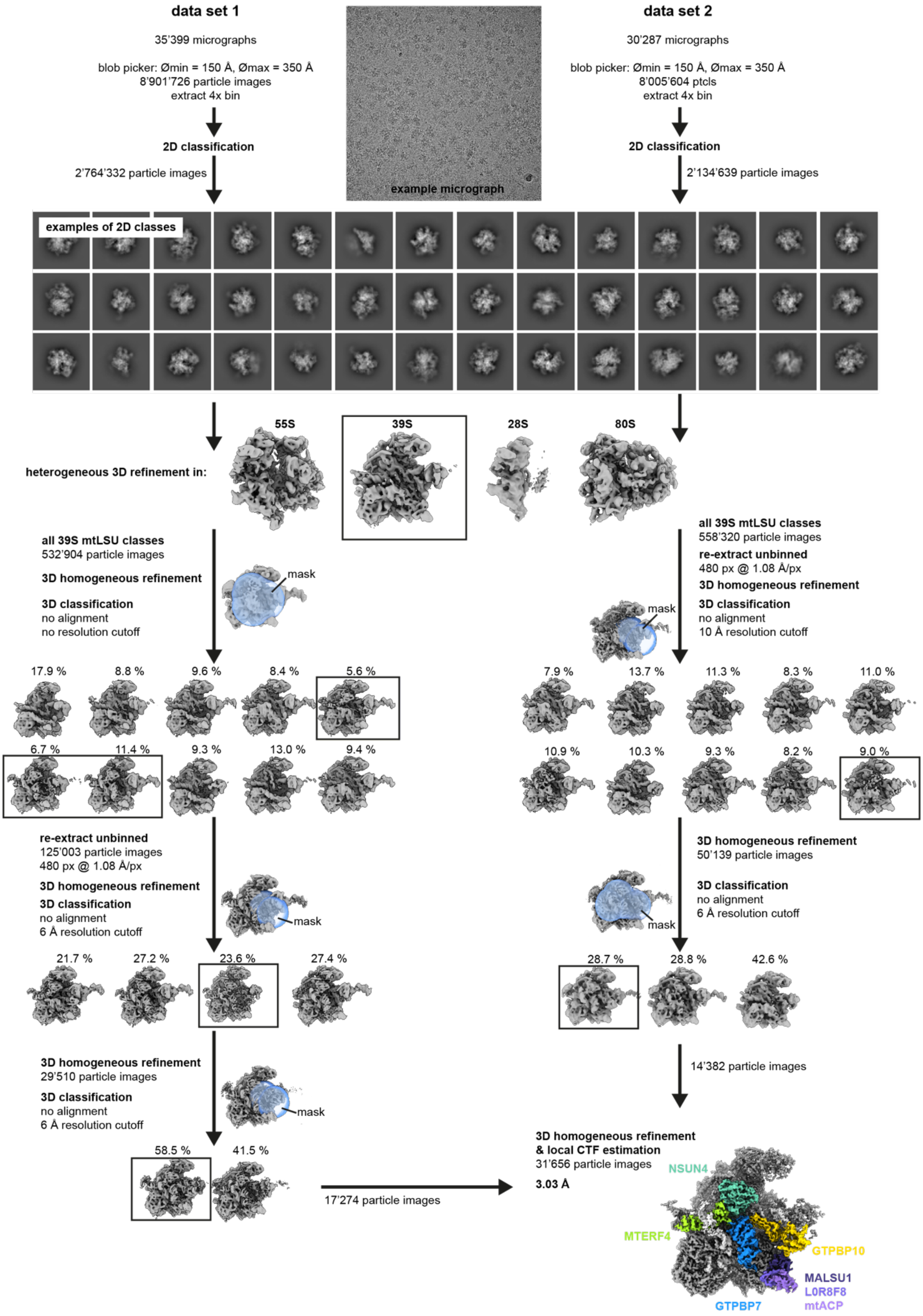


**Suppl. Fig. 5 Classification scheme for intermediate 1**

The classification scheme to derive intermediate 1 containing GTPBP10 and GTPBP7 from datasets 1 and 2 is given. An example micrograph is shown on the top. Mask that were used for local classification are shown in blue and semi-transparent. The final EM reconstruction is shown as unsharpened map and with density corresponding to the biogenesis factors colour-coded.

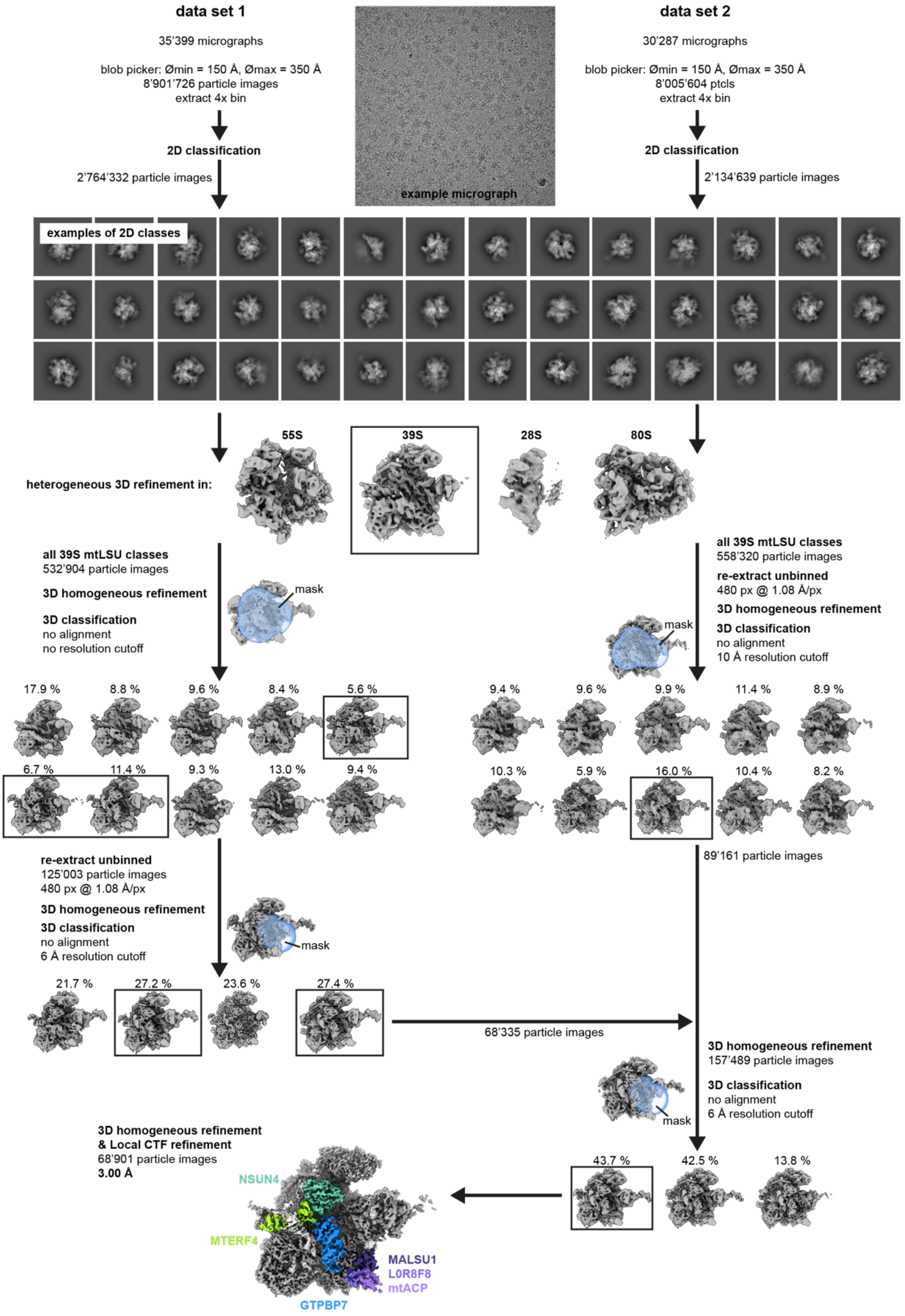


**Suppl. Fig. 6 Classification scheme for intermediate 2**

The classification scheme to derive intermediate 2 containing GTPBP7 from datasets 1 and 2 is given. An example micrograph is shown on the top. Mask that were used for local classification are shown in blue and semi-transparent. The final EM reconstruction is shown as unsharpened map and with density corresponding to the biogenesis factors colour-coded.

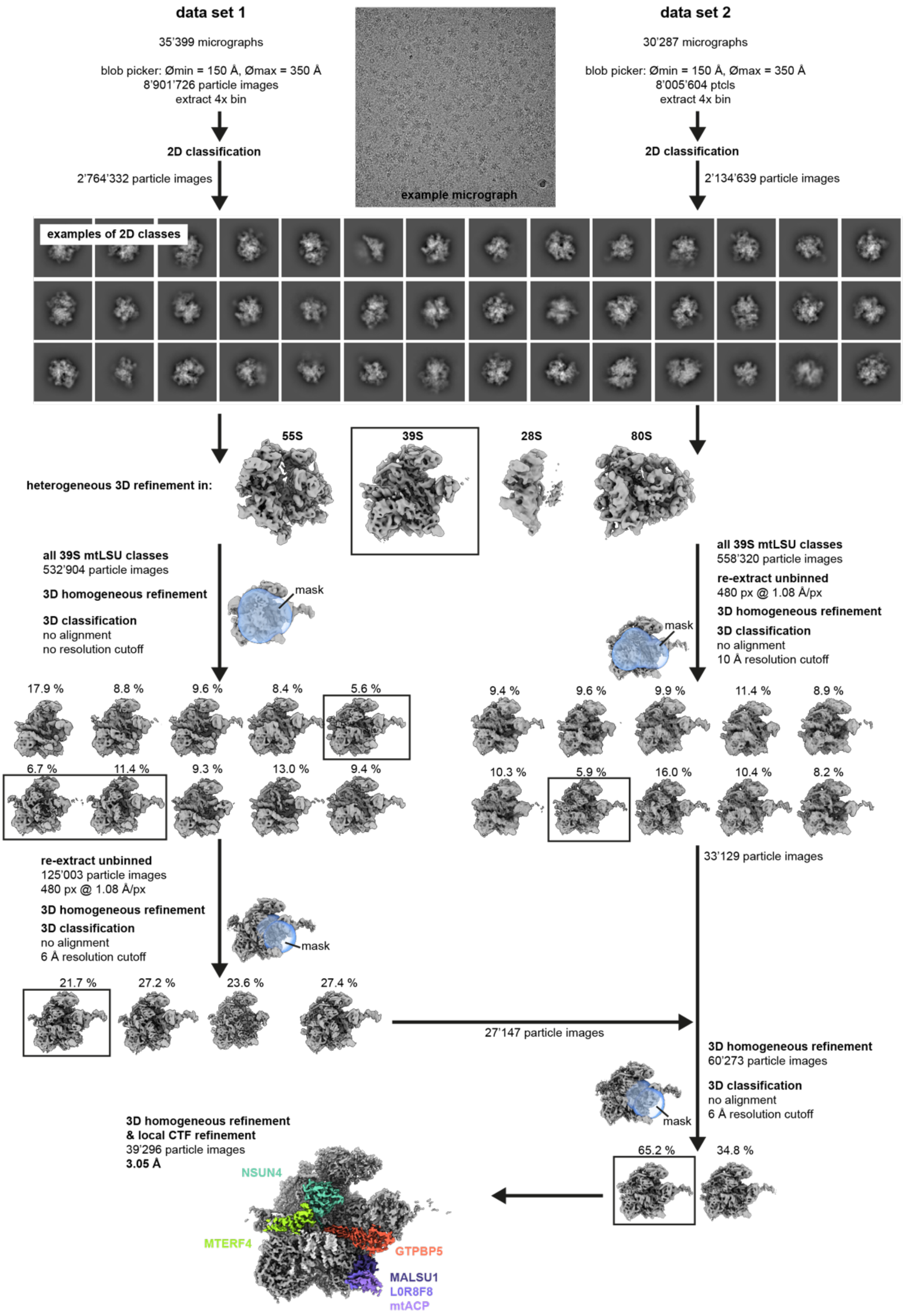


**Suppl. Fig. 7 Classification scheme for intermediate 3**

The classification scheme to derive intermediate 3 containing GTPBP5 from datasets 1 and 2 is given. An example micrograph is shown on the top. Mask that were used for local classification are shown in blue and semi-transparent. The final EM reconstruction is shown as unsharpened map and with density corresponding to the biogenesis factors colour-coded.

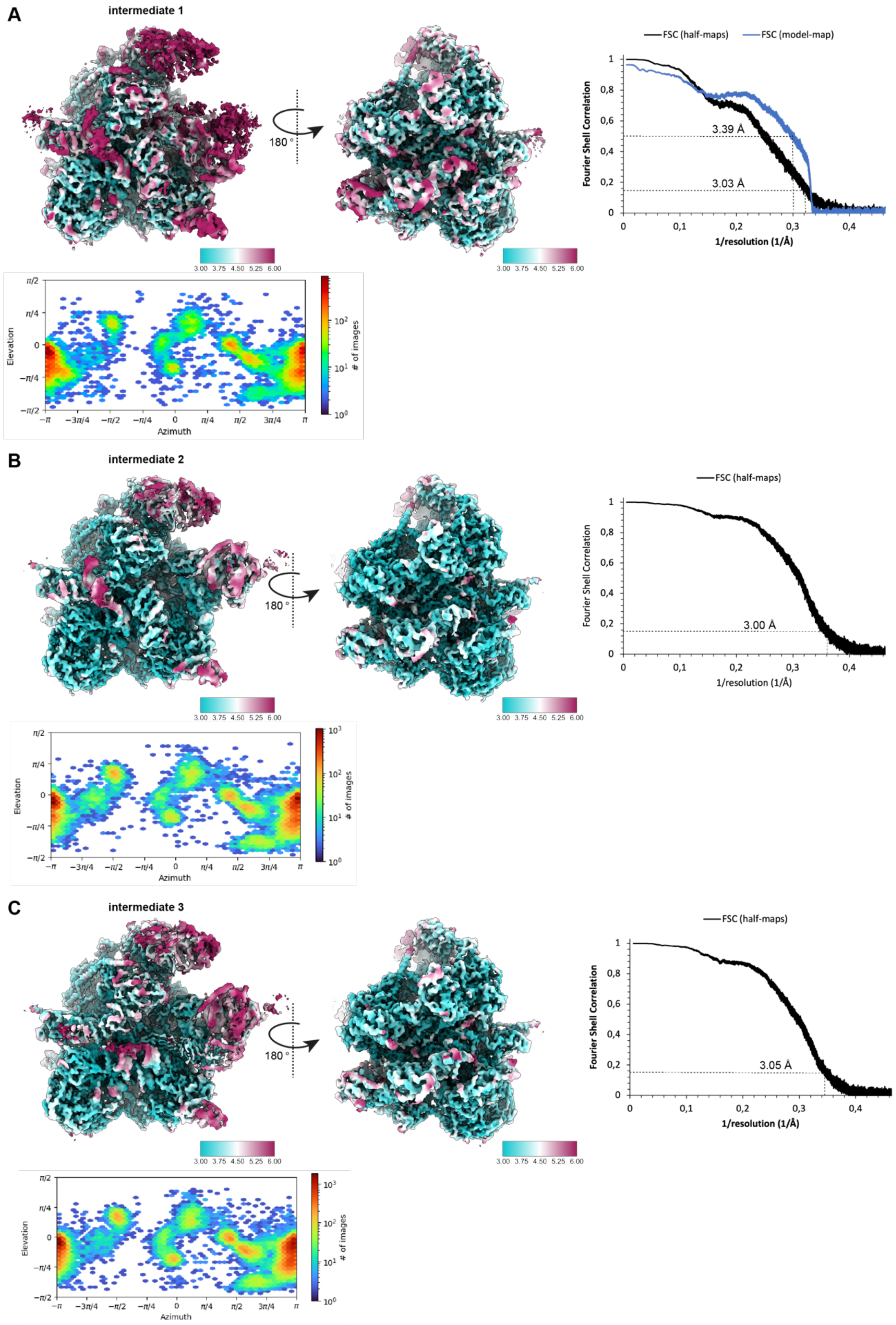


**Suppl. Fig. 8 Local resolution, FSC, and angular distribution**

Data for intermediate 1, 2 and 3 are shown in **A)**, **B)**, and **C)**, respectively. Local resolution has been estimated in cryoSPARC and is plotted according the the given colour key. The FSC curves have been calculated in PHENIX using *phenix.mtriage*. The resolution cutoff at an FSC value of 0.143 is shown via the dashed line and the masked, overall resolution value is given in Å. A 2D representation of the angular distribution of the particle images in the final reconstruction is given in form of an elevation/Azimuth heatmap (provided in radians). The number of particle images in the respective orientation is depicted as color and the corresponding color key is given on the right.
